# *Pb*AP2-FG2 and AP2R-2 function together as a transcriptional repressor complex essential for *Plasmodium* female development

**DOI:** 10.1101/2022.09.22.508975

**Authors:** Tsubasa Nishi, Izumi Kaneko, Shiroh Iwanaga, Masao Yuda

## Abstract

Gametocyte development is a critical step in the life cycle of *Plasmodium*. Despite that numbers of studies in the gametocyte development have been conducted, the molecular mechanisms regulating this process remains to be fully understood. This study investigates the functional roles of two female-specific transcriptional regulators, *Pb*AP2-FG2 and AP2R-2, in *P. berghei*. Knockout of *pbp2-fg2* or *ap2r-2* impairs female gametocyte development, resulting in developmental arrest during ookinete development. ChIP-seq analyses of these two factors indicated their colocalization on the genome, suggesting they function as a complex. These analyses also revealed that their target genes contained a variety of genes, including both male and female-enriched genes. Moreover, differential expression analyses showed that these target genes were upregulated through the disruption of *pbp2-fg2* or *ap2r-2*, indicating that these two factors function as a transcriptional repressor complex in female gametocytes. Further target analysis demonstrated a significant overlap between the target genes of *Pb*AP2-FG2 and AP2-G, suggesting that repression of early gametocyte genes activated by AP2-G is one of the key roles for this female transcriptional repressor complex. Our results indicate that the *Pb*AP2-FG2-AP2-R2 complex-mediated repression of the target genes supports the female differentiation from early gametocytes.

**Author Summary:** Gametocyte development in *Plasmodium* parasites, a causative agent of malaria, is an essential step for their transmission from vertebrate hosts to mosquitoes. Gametocytes are sexual precursor cells produced from a subpopulation of asexual blood-stage parasites. Upon uptake by mosquitoes through blood feeding, the male and female gametocytes become microgametes and macrogametes, respectively, and then they fertilize and develop into the mosquito midgut invasive stage, called ookinete. Therefore, it is crucial to understand the underlying mechanisms regulating this developmental process. This study revealed that the two female transcriptional regulators, *Pb*AP2-FG2 and AP2R-2, function together as an essential transcriptional repressor complex in *P. berghei*, the target genes of which include male, female, and early gametocyte genes activated by AP2-G. Our findings suggest that *Pb*AP2-FG2 and AP2R-2 play multiple roles in supporting the development of female gametocytes from early gametocytes.

## Introduction

*Plasmodium* parasites are the causative agent of malaria, one of the most severe infectious diseases worldwide. The spread of the parasites among individuals occur through mosquito bites, resulting in more than 200 million cases and 500 thousand deaths yearly [1]. Parasite transmission from vertebrate hosts to mosquitoes is involved in the sexual development of the parasite [2, 3]. During asexual reproduction in the host blood, subpopulations of parasites differentiate into gametocytes to prepare for transmission to mosquitoes [4, 5]. When the gametocytes are taken up by mosquitoes through blood feeding, they egress from red blood cells, form gametes, and fertilize. Fertilized cells then develop into ookinetes and invade the midgut of mosquitoes, completing the transmission [6]. Parasite transmission is a critical event in the propagation of malaria, thus understanding the molecular mechanisms regulating these developmental steps is crucial for malaria epidemiology.

In *Plasmodium* spp., sexual development is triggered by AP2-G. It is an AP2-family transcription factor that is expressed in a subpopulation of blood-stage parasites, with disruption of the gene resulting in complete loss of the parasite’s capability to produce gametocytes [7, 8]. Furthermore, it has been reported that intentional conversion of parasites into the sexual stage could be achieved by conditional induction of AP2-G in both *P. falciparum* and *P. berghei* [9, 10]. We previously conducted chromatin immunoprecipitation (ChIP) followed by high-throughput sequencing (ChIP-seq) analyses of AP2-G in *P. berghei* and identified its target genes [11]. Among these targets of AP2-G, we found several important transcription factors for sexual development. These targets include *ap2-g2*, a transcription factor that is expressed in both male and female gametocytes. AP2-G2 functions as a transcriptional repressor to ensure the alteration of cell fate from the asexual blood stage to the sexual stage [12, 13]. The target genes of AP2-G also include a female-specific transcription factor gene, *ap2-fg*. AP2-FG activates most female-specific genes, and disruption of this gene results in the formation of abnormal female gametocytes [14]. In addition to these gametocyte-specific transcription factors, the zygote transcription factor gene *ap2-z* was also identified as a target of AP2-G [15]. Therefore, it was considered that AP2-G comprehensively activates transcriptional regulator genes essential for *Plasmodium* sexual development.

*ap2-o3* and *ap2r-2* are also a target gene of AP2-G.It has been reported that in both *P. berghei* and *P. yoelii*, disruption of *ap2-o3* results in arrest of parasite development during ookinete development, thereby helping derive the name *ap2-o3* [16, 17]. AP2-R2, on the other hand, is expressed in female gametocytes, and disruption of this gene impairs ookinete development [11]. Here, we report that *Pb*AP2-O3 and AP2R-2 are expressed in female gametocytes and function together as a transcriptional repressor complex in *P. berghei*. Accordingly, we renamed *Pb*AP2-O3 *Pb*AP2-FG2. Moreover, differential expression analyses and ChIP-seq analyses revealed that *Pb*AP2-FG2 and AP2-R2 repress various genes, including female, male, and early gametocyte genes, to support female development. Recently, Li *et al*. reported that the ortholog of *Pb*AP2-FG2 in another rodent malaria parasite, *P. yoelii*, (*Py*AP2-FG2) also functions as a transcriptional repressor in female gametocytes [18]. However, the other conclusions derived from their RNA-seq and ChIP-seq data failed to corroborate our results for *P. berghei,* despite the two species being phylogenetically very close. One notable difference is their main conclusion that *Py*AP2-FG2 globally represses male-specific genes. We addressed these inconsistencies by reassessing their data and concluded that in both *P. berghei* and *P. yoelii*, the roles of AP2-FG2 in female development are identical.

## Results

### *Pb*AP2-FG2, expressed in female gametocytes, is essential for their development

*Pb*AP2-FG2 (encoded by PBANKA_1015500), previously named AP2-O3, is an AP2-family transcription factor, conserved across the *Plasmodium* species. It possesses a single AP2 domain near its N-terminus and an AP2-coincident domain mostly at the C-terminus (ACDC) domain (Fig 1A). Modrzynska *et al*. demonstrated that disruption of this gene results in the failure of the majority of zygotes to form an apical protrusion in *P. berghei* [16]. However, they did not examine the expression of *Pb*AP2-FG2 in *P. berghei*; thus, it is unclear whether *Pb*AP2-FG2 is directly involved in ookinete development. Our previous ChIP-seq data demonstrated that the upstream region of *pbap2-fg2* harbored binding sites of AP2-G and AP2-FG, which are transcriptional activators expressed in early gametocytes and female gametocytes, respectively (Fig 1B) [11, 14]. This indicates that *Pb*AP2-FG2 is possibly expressed in female gametocytes. To identify the stage at which *Pb*AP2-FG2 functions, we first generated a parasite line expressing GFP-fused *Pb*AP2-FG2 (*Pb*AP2-FG2::GFP, S1A Fig) and assessed its expression pattern. In the blood stage, *Pb*AP2-FG2 expression was observed in the nucleus of female gametocytes but not in the other stages including male gametocytes (Fig 1C). When cultured in an ookinete culture medium, *Pb*AP2-FG2::GFP parasites produced banana-shaped ookinetes, confirming that the GFP fusion did not impair *Pb*AP2-FG2 function. During ookinete development, no fluorescence was detected at any stage of the parasites (Fig 1C). These results collectively indicated that like *Py*AP2-FG2, *Pb*AP2-FG2 is only expressed in female gametocytes during sexual development and presumably functions in females instead of zygotes.

**Fig 1.**
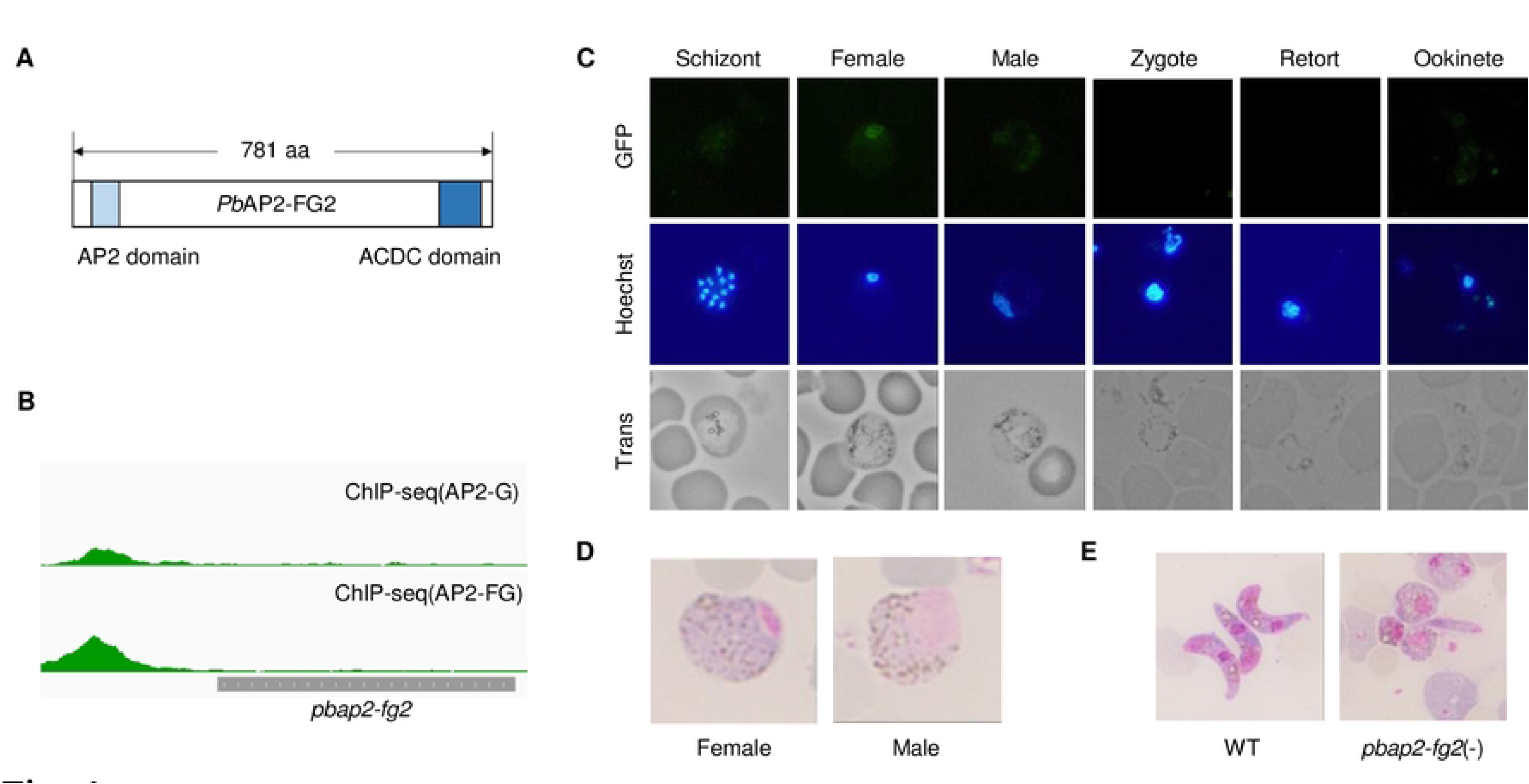
Expression pattern and knockout phenotype of *pbap2-fg2*. (A) Schematic diagram of protein features in *Pb*AP2-FG2. (B) An Integrative Genomics Viewer (IGV) image showing peaks identified in ChIP-seq analysis of AP2-G and AP2-FG in the upstream region of *pbap2-fg2*. The grey bar indicates the gene body of *pbap2-fg2*. (C) Fluorescence analysis of *Pb*AP2-FG2::GFP during blood-stage and sexual development. Nuclei were stained with Hoechst 33342. (D) Giemsa-stained images showing gametocytes of *pbap2-fg2*(-) parasites. (E) Giemsa-stained images showing ookinetes of WT and *pbap2-fg2*(-) at 20 h after starting ookinete cultures. (F) Cross-fertilization assay among *pbap2-fg2*(-), *p48/45*(-) and *p47*(-). The number of normal and abnormal ookinetes are indicated as white and grey bars, respectively.

To investigate the function of *Pb*AP2-FG2, we developed *pbap2-fg2* knockout parasites [*pbap2-fg2*(-), S1B Fig] and evaluated the phenotype in detail. *pbap2-fg2* was disrupted by double cross-over homologous recombination of an *hdhfr* expression cassette, with the transfection conducted twice to obtain two independent clonal lines. The resultant parasites formed morphologically normal female and male gametocytes (Fig 1D), and the male gametocytes showed normal exflagellation (> 30 per 100,000 red blood cells). In the ookinete culture medium, *pbap2-fg2*(-) parasites failed to produce banana-shaped ookinetes; more than half of the fertilized population stopped developing at round zygotes, and the others, at retort-form ookinetes (Fig 1E). Furthermore, both of the clones failed to infect mosquitoes through blood feeding, confirming that *pbap2-fg2*(-) parasites completely lost the ability to produce normal ookinetes. These results corroborated the findings of Modrzynska *et al.* [16]. Next, we performed a cross-fertilization assay to evaluate whether female or male gametocytes of *pbap2-fg2*(-) were capable of forming banana-shaped ookinetes upon fertilization with normal gametocytes. We observed that crossing of *pbap2-fg2*(-) with a line that produced infertile females [*p47*(-)] [19] led to no female gametocytes being converted to banana-shaped ookinetes (Fig 1F). Development of their fertilized females was arrested at the round zygote or retort-form ookinete, recapitulating the scenario of *ap2-fg2*(-) parasites cultured alone. In contrast, when *pbap2-fg2*(-) was crossed with a line that produces infertile males [*p48/45*(-)] [20], approximately 30% of female gametocytes were converted to banana-shaped ookinetes, which was approximately as much as when *p48/45*(-) and *p47*(-) were crossed (Fig 1F), demonstrating the ability of *pbap2-fg2*(-) to produce normal male gametocytes. Collectively, these results revealed that only female gametocytes were abnormal in *pbap2-fg2*(-), which, in turn, affected their ookinete development. Together with the fluorescence analysis, these results strongly suggested that *Pb*AP2-FG2 is essential for the development of normal female gametocytes, which is consistent with the study of the *Py*AP2-FG2 reported by Li *et al*. [18].

### Disruption of *pbap2-fg2* affected the female transcriptome

To further investigate the effect of disrupting *pbap2-fg2* on female development, we performed RNA-seq analysis on gametocyte-enriched populations of wild-type ANKA strain (WT) and *pbap2-fg2*(-), and compared their transcriptomes. The total RNA was harvested from parasites enriched with gametocytes, which were prepared by killing asexual parasites with sulfadiazine treatment, and then sequenced using next-generation sequencing. Differentially expressed genes (DEGs) between WT and *pbap2-fg2*(-) were identified by analyzing the sequence data using DESeq2 after excluding genes with reads per kilobase of transcript per million mapped reads (RPKM) < 10 as the minimum threshold (S1A Table). In *pbap2-fg2*(-) parasites, 180 genes were significantly downregulated [log_2_(fold change) < -1, *p*-value adjusted for multiple testing with the Benjamini-Hochberg procedure (*p*-value adj) < 0.05], and 96 genes were significantly upregulated [log_2_(fold change) > 1, *p*-value adj < 0.05] compared to the WT (Fig 2A, S1B and S1C Table). To evaluate how disruption of *pbap2-fg2* affects gametocyte transcriptome, we assessed the expression of these DEGs in previously reported sex-specific RNA-seq data [21]. We identified genes more than fourfold enriched with *p*-value adj < 0.001 in each sexual stage compared to the other and asexual blood stages as sex-enriched genes and obtained 504 female-enriched genes and 438 male-enriched genes (S2A and S2B Table). The genes downregulated in *pbap2-fg2*(-) contained 54 female-enriched genes and nine male-enriched genes, clearly highlighting the dominance of female-enriched genes (Fig 2A and 2B). Furthermore, overall, log_2_(fold change) of female-enriched genes tended to be lower than the other genes (*p*-value = 1.4 × 10^-18^ by two-tailed Student’s t-test). In contrast, the upregulated genes showed no specific enrichment in the female- or male-enriched genes (Fig 2A and 2C). These results indicated that disruption of *pbap2-fg2* impaired the female transcriptome, causing downregulation of female-enriched genes.

**Fig 2.**
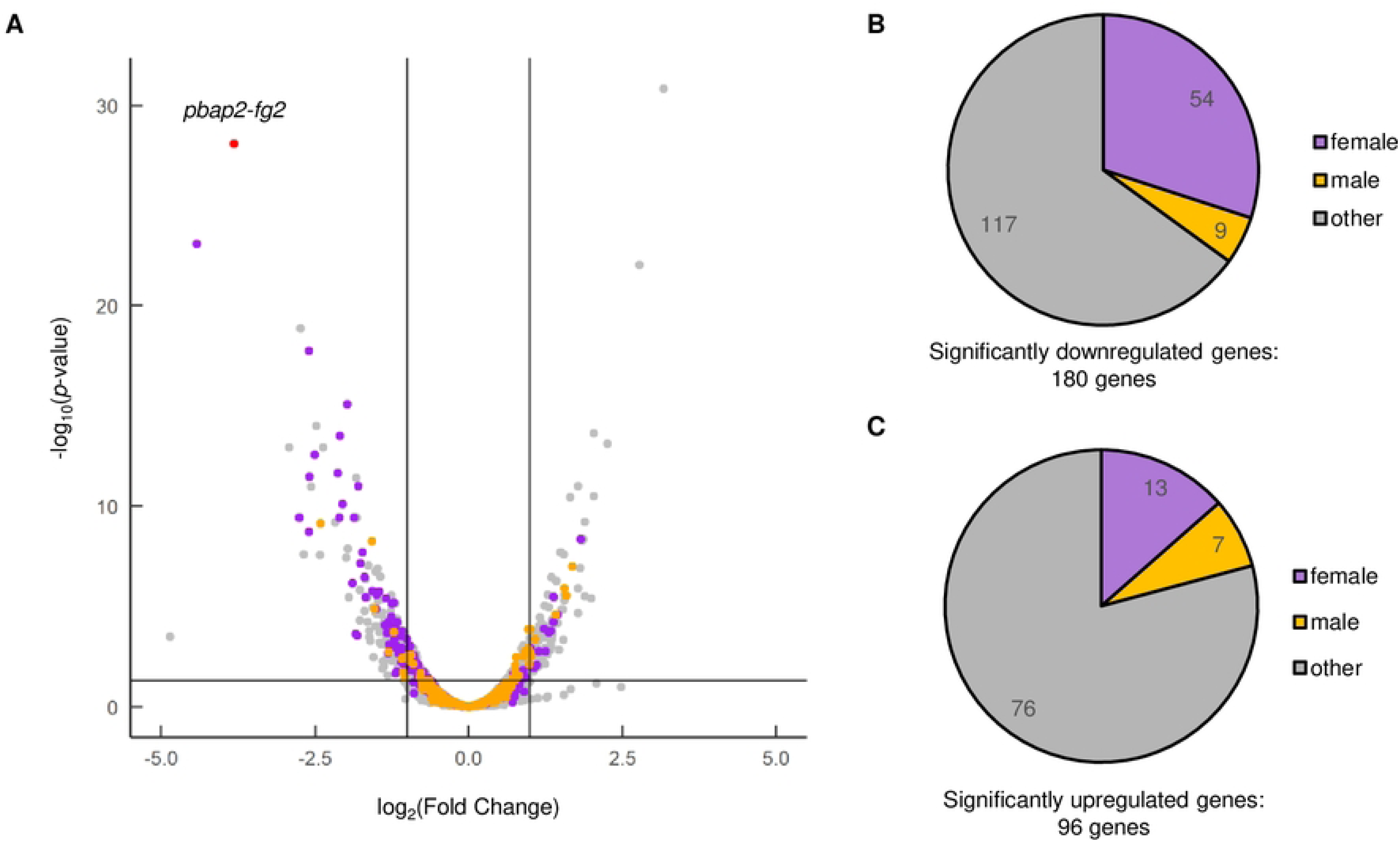
Differential expression analysis between WT and *pbap2-fg2*(-). (A) A volcano plot showing differential expression of genes in *pbap2-fg2*(-) compared to WT. Purple and orange dots represent female and male-enriched genes, respectively. A red dot indicates *pbap2-fg2*. A horizontal line indicates *p*-value of 0.05, and two vertical lines indicate log_2_(Fold Change) of -1 and 1. (B) Classification of significantly downregulated genes into sexual stage-enriched gene sets. Genes not enriched in either “female” or “male” are classified as “other”. (C) Classification of significantly upregulated genes into sexual stage-enriched gene sets.

### *Pb*AP2-FG2 targets a wide variety of genes, binding to specific **sequences**

Differential expression analysis between WT and *pbap2-fg2*(-) suggested that *Pb*AP2-FG2 was involved in transcriptional regulation in female gametocytes. Therefore, we employed ChIP-seq analysis to identify the binding motif of *Pb*AP2-FG2 and its target genes. We performed ChIP experiments with *Pb*AP2-FG2::GFP using an anti-GFP antibody, followed by the sequencing of the DNA fragments purified from the immunoprecipitated chromatin and the input cell lysate. From the sequence data, peaks were called with fold enrichment > 3.0 and *q*-value < 0.01 using the macs2 program, setting the sequence data of input DNA as a control. Two biologically independent experiments were performed, the results of which were comparable based on the genome-wide peak pattern of the data (Fig 3A). We identified 1321 and 1648 peaks in Experiments 1 and 2, respectively, and the locations of 1231 peaks (93.2% of the peaks from Experiment 1) overlapped between the two experiments, suggesting that the data had high reproducibility. To further evaluate the reproducibility of each peak, IDR1D analysis was performed on the two data [22, 23]. In this analysis, peaks were ranked according to their *p*-values within each replicate, and the ranks were compared between the two experiments. According to the consistency of ranks across the two experiments, the irreproducible discovery rate (IDR) score, which defines the reproducibility of each peak, was calculated for each one. As the ranks of each peak lose consistency between the replicates, the peaks have higher IDR scores; hence, peaks with small IDR scores are considered reliable. The results depicted that 638 peaks had an IDR < 0.01 (Fig 3B, S3A and S3B Table), and we decided to utilize these peaks for further analysis as they are highly reproducible.

**Fig 3.**
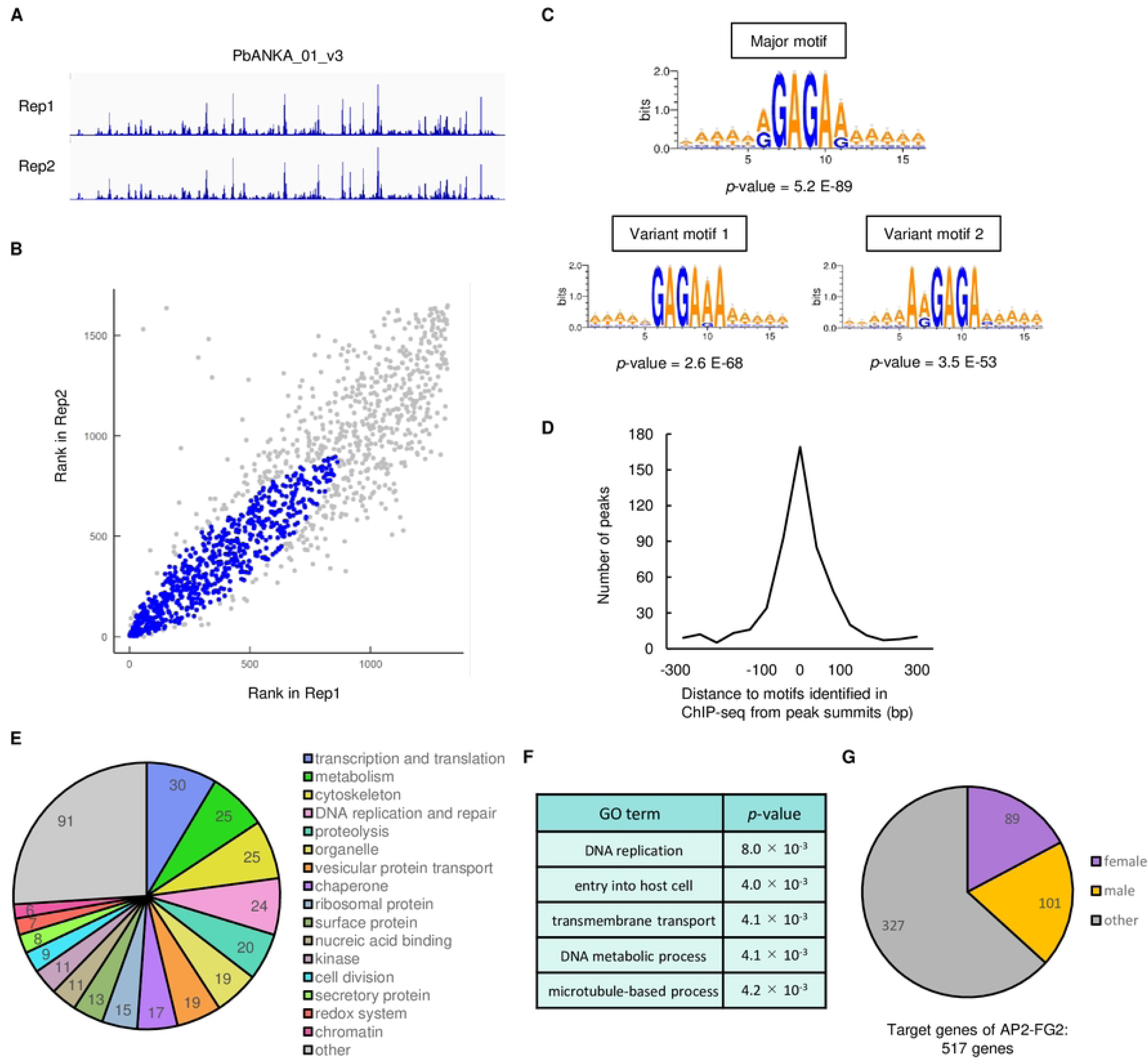
ChIP-seq analysis using *Pb*AP2-FG2::GFP. (A) IGV images showing peaks identified in the ChIP-seq experiment 1 and 2 of *Pb*AP2-FG2 on the chromosome 1. (B) IDR1D analysis between the ChIP-seq experiment 1 and 2. The rank of peaks according to their *p*-value for each experiment is plotted against each other. Peaks with IDR < 0.01 are indicated as blue dots. (C) Motifs enriched within 50 bp from peak summits identified in the ChIP-seq of *Pb*AP2-FG2. The logos were depicted using WebLogo 3 (http://weblogo.threeplusone.com/). (D) Distance between peak summits and the nearest major or variant motifs. (E) Classification of target genes into 17 groups according to their functional annotation. (F) Gene ontology analysis for target genes of *Pb*AP2-FG2. Terms with *p*-value < 0.01 are shown. (G) Classification of target genes of *Pb*AP2-FG2 into sexual stage-enriched gene sets.

We first attempted to identify the binding motif of *Pb*AP2-FG2 by searching for statistically enriched sequences around the highly reproducible peaks. We searched for 6-bp motifs on sequences within 100 bp from each summit and found enrichment of several motifs using Fisher’s exact test. These motifs were unified, and RGAGAR (R = A or G) was identified as the most significantly enriched motif in the peak regions with a *p*-value of 5.2 × 10^-89^ (Fig 3C). In addition to the RGAGAR motifs, we found GAGARA and ARGAGA as enriched motifs with a *p*-value of 2.6 × 10^-68^ and 3.5 × 10^-53^, respectively (Fig 3C). These motifs appeared to be variants of the most enriched motif, sharing GAGA within their sequences. Accordingly, we hereafter refer to the RGAGAR motif as the major motif and the GAGARA and ARGAGA motifs as the variant motifs 1 and 2, respectively. Searching for these three motifs around peak summits revealed that 85% of the peaks had at least one of these enriched motifs within 300 bp of the summit. Moreover, for more than half of these peaks, the distance between the peak summit and the nearest motif was within 50 bp (Fig 3D). These results indicated that *Pb*AP2-FG2 binds to the major motif RGAGAR and its variant motifs.

Next, we analyzed the genomic location of the peaks identified by ChIP-seq analysis to determine the potential targets of *Pb*AP2-FG2. The analysis revealed that *Pb*AP2-FG2 bound to the upstream regions (within 1200 bp from ATG) of 517 genes (S3C Table). Of the 517 target genes, 350 have been functionally annotated on PlasmoDB (https://plasmodb.org). We classified these 350 genes into functional groups to evaluate functional characteristics of the target genes (Fig 3E and S3C Table). The target genes contained some groups of genes seemingly expressed in female gametocytes, such as “cytoskeleton” and “secretory protein”. The group “cytoskeleton” had some genes encoding inner membrane complex and myosin proteins [24, 25], and the group “secretory protein” contained *warp* and some secreted ookinete protein genes [26, 27]. Meanwhile, some other functional groups, *such as* “DNA replication and repair” and “cell division”, did not seem to be related to female development. To further investigate whether genes of any specific function were enriched in the targets, we performed a gene ontology (GO) analysis. The GO analysis revealed that the target genes were most enriched in the term “DNA replication,” which included putative DNA replication licensing factor genes, DNA polymerase subunit genes and so on, with *p*-value of 8.0 × 10^-4^ (Fig 3F). In addition, genes that belong to the GO terms “entry into host cell,” “transmembrane transport,” “DNA metabolic process,” and “microtubule-based process” were also found to be enriched (*p*-value < 0.01) (Fig 3F).

Next, we evaluated the composition of sex-enriched genes among the targets of *Pb*AP2-FG2. Of the 517 target genes, 90 were female-enriched (Fig 3G, S3C Table). However, in these female-enriched target genes, genes of any specific function did not appear enriched. The targets also contained 101 male-enriched genes (Fig 3G, S3C Table), including most of the genes that were classified into the functional groups “DNA replication and repair” and “cell division”. Because *Pb*AP2-FG2 is a female-specific transcription factor, the number of male-enriched genes in the targets seemed too large, again implying the possible role of *Pb*AP2-FG2 as a transcriptional repressor. Concordantly, when the association between the target genes of *Pb*AP2-FG2 and genes downregulated in *pbap2-fg2*(-) was assessed, only four target genes were significantly downregulated. Therefore, we considered that the downregulation of genes in *pbap2-fg2*(-) was not a direct effect of disrupting *pbap2-fg2*.

### Target genes of *Pb*AP2-FG2 were upregulated in *ap2-fg2*(-)

To evaluate how the disruption of *Pb*AP2-FG2 affected the transcription of its target genes, we compared the target genes and DEGs identified in the RNA-seq analysis. Intriguingly, the target genes were enriched in the genes significantly upregulated in *pbap2-fg2*(-) (40 of the 96 upregulated genes, *p*-value = 6.9 × 10^-15^ by Fisher’s exact test), while only four targets were included in the significantly downregulated genes, as mentioned above (Fig 4A, S3C Table). In addition, although not more than 2-fold, the other 47 target genes were upregulated, with a *p*-value adj < 0.05. Furthermore, comparing the log_2_(fold change) distribution of the target genes with that of the other genes revealed that the target genes tended to be upregulated in *pbap2-fg2*(-) with a *p*-value of 8.2 × 10^-58^ by two-tailed Student’s t-test (Fig 4A). Therefore, we considered that *Pb*AP2-FG2 repressed the transcription of its target genes in female gametocytes. An upregulation pattern of the target genes was also observed in female and male-enriched genes with a *p*-value of 3.9 × 10^-13^ and 1.2 × 10^-4^ by two-tailed Student’s t-test, respectively (Fig 4B and C), indicating a role of *Pb*AP2-FG2 in repressing its target genes regardless of their expression specificity.

**Fig 4.**
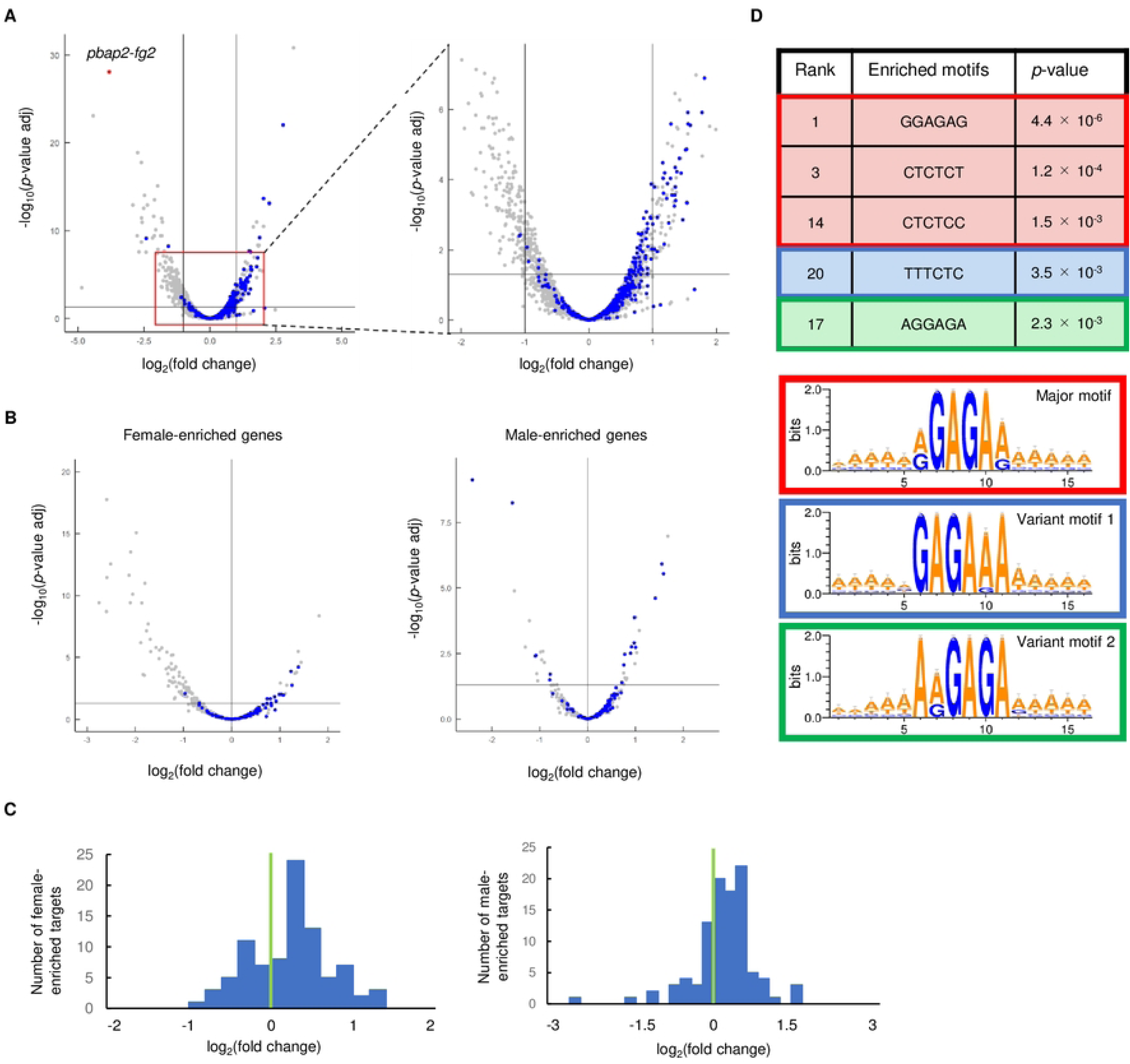
Relationship between target genes of *Pb*AP2-FG2 and DEGs in *pbap2-fg2*(-). (A) A volcano plot showing DEGs in *pbap2-fg2*(-) compared to WT. Blue dots represent the target genes of *Pb*AP2-FG2, and a red dot indicates *pbap2-fg2*. A horizontal line indicates *p*-value of 0.05, and two vertical lines indicate log_2_(Fold Change) of -1 and 1. All genes with RPKM > 10 are depicted in the left panel. For the right panel, region from log_2_(Fold Change) of -2 to 2 and from -log_10_(*p*-value) of 0 to 7.5 is magnified. (B) Volcano plots showing DEGs in *pbap2-fg2*(-) for female and male-enriched genes (the left and right panel, respectively). Blue dots represent the target genes of *Pb*AP2-FG2. A horizontal line indicates a *p*-value of 0.05, and a vertical line indicates log_2_(Fold Change) of 0. (C) Distribution of log_2_(Fold Change) values for female and male-enriched target genes of *Pb*AP2-FG2 (the left and right graph, respectively). A green line indicates log_2_(Fold Change) of 0. (D) Six-bp DNA motifs enriched within the upstream region (300 to 1200 bp from ATG) of genes upregulated in *pbap2-fg2*(-). The major and variant motifs are each indicated in different color boxes. The ranks were assigned to all enriched motifs according to their *p*-values.

Considering the above results, we examined whether the binding motifs of *Pb*AP2-FG2 were enriched in the upstream region (300–1200 bp from ATG) of the upregulated genes compared to that of the other genes. Through this analysis, we found that one of the motifs that belong to the major motifs, GGAGAG, was found to be the most enriched by Fisher’s exact test (*p*-value = 4.4 × 10^-6^, Fig. 4D). Additionally, two other major motifs (CTCTCT and CTCTCC) and one of the variant motifs 1 and 2 (TTTCTC and AGGAGA, respectively) were also found to be enriched, with a *p*-value < 0.005 (Fig 4B). These results strongly suggested that the upregulation of genes in *pbap2-fg2*(-) was primarily a direct effect of its disruption.

### The binding motifs of *Pb*AP2-FG2 functioned as a *cis*-acting repressive element

To confirm that *Pb*AP2-FG2 functions as a transcriptional repressor upon binding to the major and variant motifs, we evaluated the effects of disrupting motifs in the upstream region of target genes. First, we generated parasites expressing GFP-fused *Pb*AP2-FG2 by the CRISPR/Cas9 system (*Pb*AP2-FG2::GFP^C^, S1C Fig) using Cas9-expressing parasites called Pbcas9 [28]. After developing *Pb*AP2-FG2::GFP^C^, we introduced point mutations in the motif upstream of *psh3*. Parasite-specific helicase 3 (PSH3) is a helicase conserved in apicomplexan parasites, which in *P. falciparum* is reported to be expressed in trophozoites and schizonts and is essential for asexual stage development [29]. The female and male gametocyte RNA-seq data indicated that *psh3* is transcribed in male but not female gametocytes. Using the CRISPR/Cas9 system, we changed the motif in the peak region upstream of *psh3* from GGAGAA to atAtAt (Fig 5A).

**Fig 5.**
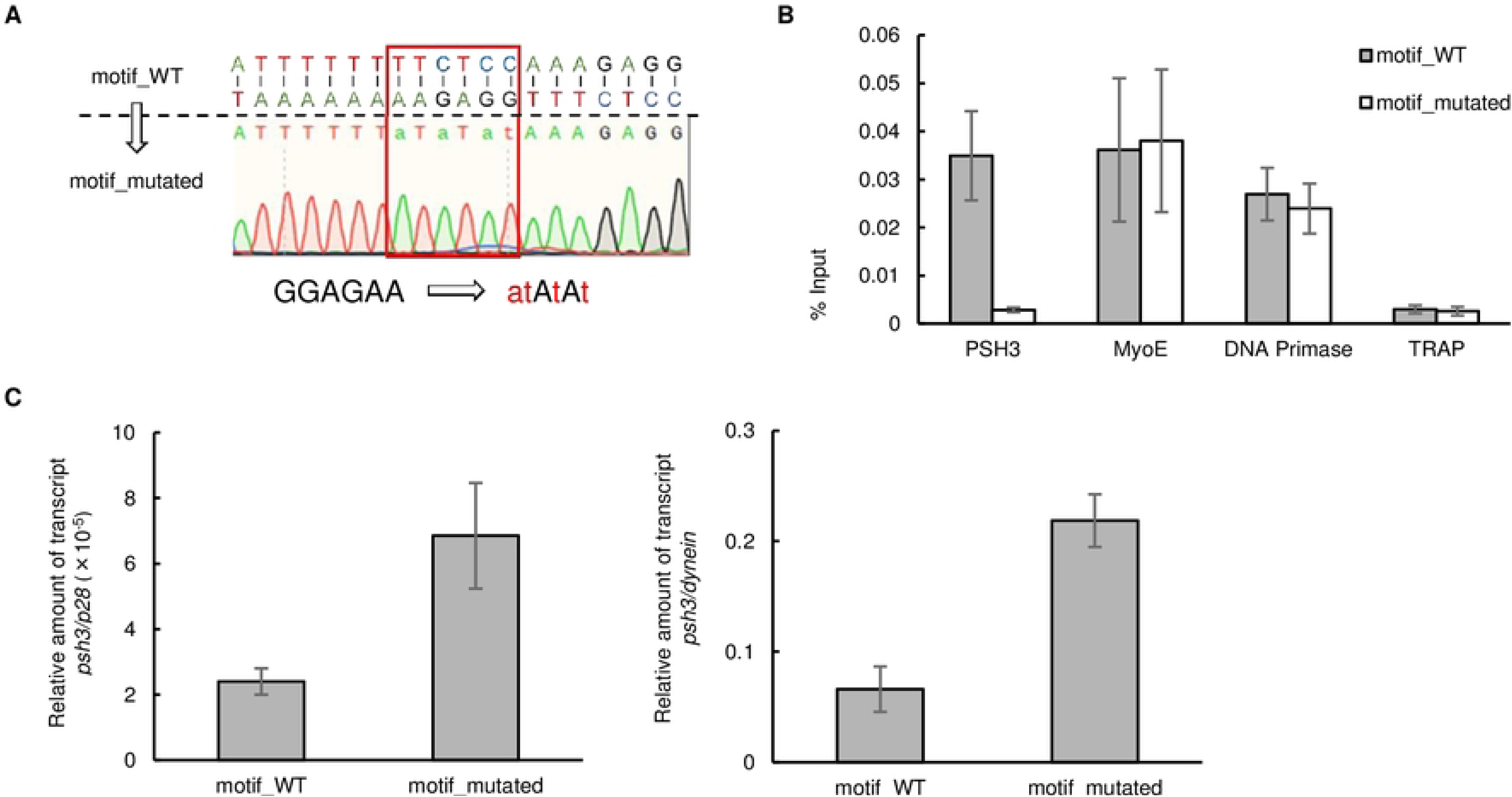
Disruption of the binding motif of *Pb*AP2-FG2 in the upstream region of *psh3*. (A) Genomic sequence around the binding motif of *Pb*AP2-FG2 located upstream of *psh3*, and Sanger sequence result of the region in motif_mutated parasites. (B) ChIP-qPCR analysis of *Pb*AP2-FG2 at the mutated site upstream of *psh3*. Grey and white bars indicate %input for motif_WT and motif_mutated, respectively. Error bars indicate the standard error of the mean %input values from three independent experiments. [PSH3: Parasite Specific Helicase 3, MyoE: Myosin E, DNA Primase: DNA Primase large subunit, TRAP: Thrombospondin-Related Anonymous Protein] (C) RT-qPCR analysis of *psh3* in motif_WT and motif_mutated parasites. The relative transcript level of *psh3* against *p28* is presented. Error bars indicate the standard error of the mean from three independent experiments.

To evaluate the binding of *Pb*AP2-FG2 at the mutated site, we first performed ChIP coupled with quantitative PCR (ChIP-qPCR) analysis using *Pb*AP2-FG2::GFP^C^ with the wild-type (motif_WT) or mutated (motif_mutated) motif upstream of *psh3*. In the motif_mutated parasites, the amount of immunoprecipitated DNA fragments relative to input DNA (%input value) was significantly decreased at the mutated site compared to the motif_WT. In contrast, at the other sites, the upstream region of *myoE* and the DNA primase large subunit gene as positive controls and *trap* as a negative control, the %input values were comparable between the motif_WT and motif_mutated parasites (Fig 5B). Together with the ChIP-seq results, these results strongly indicated that the ChIP-seq-identified motifs are the binding motifs of *Pb*AP2-FG2.

Subsequently, using these mutants, we assessed how the binding of *Pb*AP2-FG2 affects downstream transcription by reverse transcription quantitative PCR (RT-qPCR) analysis. Total RNA was harvested from motif_WT and motif_mutated parasites treated with sulfadiazine, and the relative amount of *psh3* transcripts to *p28* transcripts was analyzed. The results demonstrated that the relative transcript level of *psh3* in motif_mutated parasites was more than 2.5-fold higher than in motif_WT (Fig 5C, left graph). Since *psh3* is usually expressed in male gametocytes but not in females, we supposed that it is essential to exclude the possibility that the male-to-female ratio affected the result. Accordingly, we evaluated the amount of *psh3* transcripts relative to that of a male-specific gene, the dynein heavy chain gene (PBANKA_0416100). The result was comparable to when *p28* was used as a control; the relative transcript level of *psh3* was more than 3-fold higher in motif_mutated than in motif_WT (Fig 5C, right graph). Collectively, these results indicated that the major motif functions as a *cis*-regulatory element for repressing downstream genes.

### *Pb*AP2-FG2 requires a co-repressor AP2R-2 to repress its target genes

We preiously identified two putative transcriptional regulator genes, *ap2r-1* and *ap2r-2*, as a target gene of AP2-G and AP2-FG [11]. Of these, *ap2r-1* functions as a transcriptional activator in zygotes and is renamed *ap2-z* [15], but the functional role of *ap2r-2* remains unknown. AP2R-2 has an ACDC domain at its C-terminus, but no AP2 domain. It is expressed in females, and *ap2r-2* knockout parasites [*ap2r-2*(-)] are not able to form banana-shaped ookinetes [11]. Therefore, we hypothesized that AP2R-2 might play an essential role in female development and assessed its function. First, we developed a parasite line expressing GFP-fused AP2R-2 using CRISPR/Cas9 (AP2R-2::GFP^C^, S1D Fig). The female gametocytes of AP2R-2::GFP^C^ showed nuclear-localized fluorescence, consistent with our previous study; therefore, we performed ChIP-seq analysis at the gametocyte stage. Through this analysis, we obtained 944 highly reproducible peaks (IDR < 0.01) from 1597 and 1459 peaks identified in Experiments 1 and 2, respectively (S4A and S4B Table). Intriguingly, the genome-wide peak pattern for ChIP-seq of *Pb*AP2-FG2 and AP2R-2 seemed almost identical (Fig 6A). To evaluate the consistency of the ChIP peak pattern over the whole genome, we assessed the read coverage in the ChIP-seq data of *Pb*AP2-FG2 at the peak summits identified in the ChIP-seq of AP2R-2 and *vice versa*. The results depicted that read coverage for *Pb*AP2-FG2 was enriched at the AP2R-2 peaks, and higher fold enrichment of AP2R-2 peaks correlated with higher read count detected in the *Pb*AP2-FG2 ChIP-seq. (Fig 6B). The same was true for the read coverage of the AP2R-2 ChIP-seq at the *Pb*AP2-FG2 peaks, indicating that the peak patterns in the ChIP-seq of *Pb*AP2-FG2 and AP2R-2 were identical. Consistently, motif enrichment analysis by Fisher’s exact test revealed that the major and two variant motifs were all enriched within 100 bp from the summits of peaks identified in the ChIP-seq of AP2R-2 (*p*-value = 7.3 × 10^-151^, 2.1 × 10^-116^ and 2.7 × 10^-76^ for the major motif, variant motifs 1 and 2, respectively, Fig 6C). Collectively, these results suggested that AP2R-2 colocalizes with *Pb*AP2-FG2 on the genome and may cooperatively work with *Pb*AP2-FG2 during female development.

**Fig 6.**
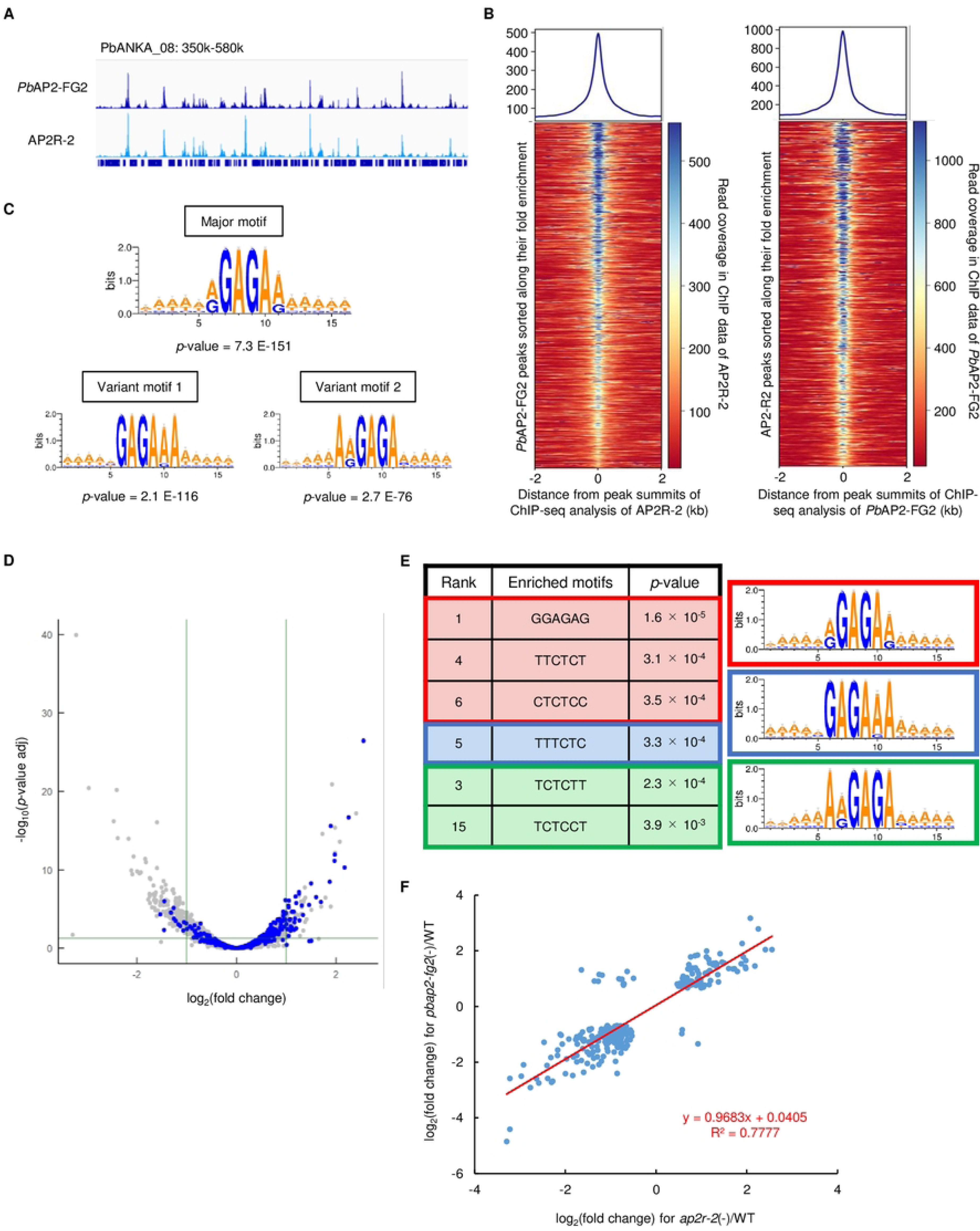
ChIP-seq of AP2R-2 and differential expression analysis between WT and *ap2r-2*(-). (A) IGV images showing peaks identified in the ChIP-seq analysis of *Pb*AP2-FG2 and AP2R-2. (B) Heat maps showing coverage in ChIP-seq of *Pb*AP2-FG2 at AP2R-2 peaks (left) and coverage in ChIP-seq of AP2R-2 at *Pb*AP2-FG2 peaks (right). Peak regions are aligned in ascending order of their fold enrichment value. Graphs on top of the heat maps show the mean coverage of all peak regions. (C) Motifs enriched within 50 bp from peak summits identified in the ChIP-seq of AP2R-2. The logos were depicted using WebLogo 3. (D) A volcano plot showing DEGs in *ap2r-2*(-) compared to WT. Blue dots represent the target genes of *Pb*AP2-FG2. A horizontal line indicates a *p*-value of 0.05, and two vertical lines indicate log_2_(Fold Change) of -1 and 1. (E) Six-bp DNA motifs enriched within upstream region of genes upregulated in *ap2r-2*(-). (F) A scatter plot showing relationship between log_2_(Fold Change) in *pbap2-fg2*(-) and *ap2r-2*(-) for genes with *p*-value < 0.05. A red line indicates a linear approximation of the plots.

To evaluate the functions of AP2R-2 in the expression of its targets, we performed differential expression analysis by comparing WT and *ap2r-2*(-). We identified 95 significantly upregulated and 222 significantly downregulated genes (Fig 6D, S5A, S5B, and S5C Table). Among the upregulated genes, 36 targets of *Pb*AP2-FG2 were detected, and overall, the target genes were tended to be upregulated in *ap2r-2*(-) with a *p*-value of 1.6 × 10^-53^ by two-tailed Student’s t-test (Fig. 6D). This upregulation tendency was also detected within female- and male-enriched genes, with a *p*-value of 5.6 × 10^-5^ and 4.2 × 10^-8^, respectively. Moreover, in the upstream region of the genes upregulated in *ap2r-2*(-), one of the major motifs, GGAGAG, was most enriched with a *p*-value of 1.6 × 10^-5^ by Fisher’s exact test, and some other binding motifs of *Pb*AP2-FG2/AP2R-2 were also significantly enriched (Fig 6E). Therefore, we considered that AP2R-2 functions as a transcriptional repressive factor in female gametocytes. The upregulated and downregulated genes identified in *pbap2-fg2*(-) and *ap2r-2*(-) presented a significant overlap with a *p*-value of 3.4 × 10^-37^ and 4.1 × 10^-62^ by Fisher’s exact test, respectively. To further evaluate the relationship between *pbap2-fg2*(-) and *ap2r-2*(-) in detail, we plotted log_2_(fold change) values in *pbap2-fg2*(-) vs. WT against those in *ap2r-2*(-) vs. WT for genes with a *p*-value less than 0.05, in both experiments (273 genes). We then constructed a linear approximation of the plots, which depicted a line with a slope of approximately 1 and a mean square correlation coefficient of 0.78 (Fig 6F). This result indicated that the fold change in the expression level for each gene was mostly comparable between *pbap2-fg2*(-) and *ap2r-2*(-); hence, disruption of *pbap2-fg2* and *ap2r-2* had almost identical effects on the female transcriptome. Therefore, we considered that AP2R-2 functions as an essential co-repressor of *Pb*AP2-FG2 in female gametocytes.

### *Pb*AP2-FG2 and AP2R-2 repress the target genes of AP2-G

In previous studies, we observed the expression patterns of AP2-G and AP2-FG and found that the expression of AP2-FG begins during the period when the expression of AP2-G decreases [11, 14]. This result indicates that a major transcriptional activator changes as early gametocytes develop into females. Considering this scenario, we hypothesized that the *Pb*AP2-FG2-AP2R-2 complex might support stage conversion from early gametocytes to female gametocytes by repressing early gametocyte genes activated by AP2-G. To address this possibility, we assessed whether the target genes of *Pb*AP2-FG2 include those of AP2-G. Consistent with our hypothesis, we found a significant overlap (105 genes) between the target genes of AP2-G and *Pb*AP2-FG2 with a *p*-value of 4.5 × 10^-7^ by Fisher’s exact test (Fig. 7A).

**Fig 7.**
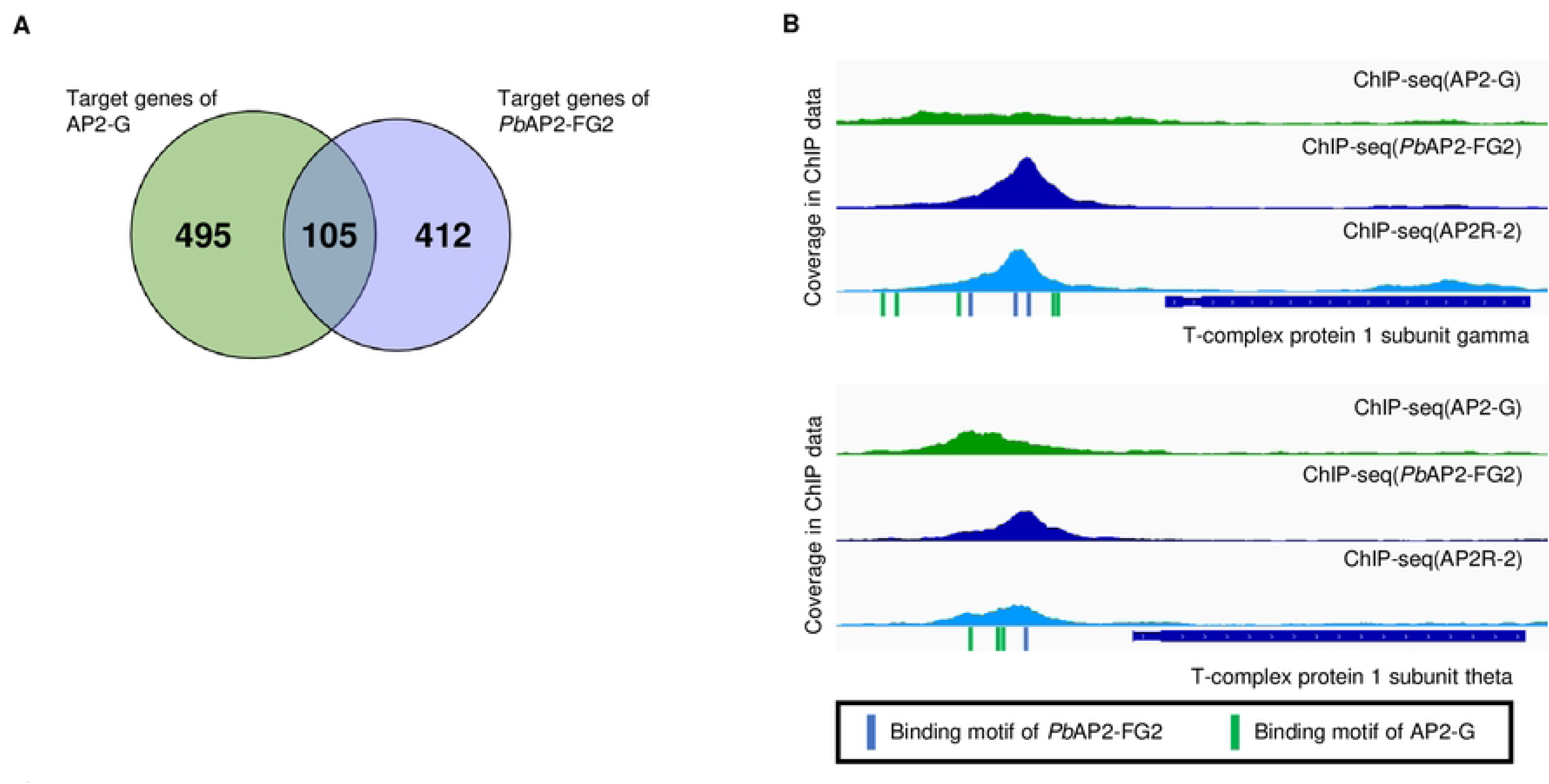
Relationship between target genes of AP2-G and *Pb*AP2-FG2. (A) A Venn diagram showing the overlap between target genes of AP2-G and *Pb*AP2-FG2. (B) IGV images showing representative peaks of ChIP-seq for AP2-G, *Pb*AP2-FG2, and AP2R-2 on the upstream of target genes common for AP2-G and *Pb*AP2-FG2. Coverage for the ChIP data is shown. Positions of the AP2-G and *Pb*AP2-FG2 binding motifs are indicated in green and blue, respectively.

Most of these common target genes between *Pb*AP2-FG2 and AP2-G have not been assessed for their function during *Plasmodium* gametocyte development. In the common target genes, we found T-complex protein 1 subunit (TCP-1) genes (Fig. 7B, S3C Table). *Plasmodium* parasites possess eight TCP-1 genes, all of which are a target of AP2-G. Although the target genes of *Pb*AP2-FG2 contained only four of these genes, peaks with fold enrichment > 2.5 were found upstream of three other TCP-1 genes in the ChIP-seq of *Pb*AP2-FG2. TCP-1s comprise a type 2 chaperonin, tailless complex polypeptide 1 ring complex (TRiC), which has been indicated to play essential roles in folding diverse polypeptides, including actin and tubulin [30–33]. In *P. falciparum*, it has been reported that TRiC is essential for the asexual blood-stage development [34, 35]. Thus, this complex is presumed to be widely required in the *Plasmodium* life cycle, including early gametocyte development, but not during female development. Notably, the common target genes of *Pb*AP2-FG2 and AP2-G also included the actin I gene, alpha-tubulin 2 gene, and a putative tubulin beta chain gene (S3C Table). These findings suggested the possibility of cytoskeletal development in female gametocytes being mostly completed during early gametocyte development.

### The function of *Py*AP2-FG2 is identical to that of *Pb*AP2-FG2

The present study revealed that transcriptional repression by the *Pb*AP2-FG2-AP2R-2 complex is essential for regulating the female transcriptome. Recently, Li *et al*. reported that *Py*AP2-FG2 is also a transcriptional repressor in female gametocytes, but their other conclusions differed from what we have revealed here [18]. The first is regarding the role of AP2-FG2, which was primarily based on their RNA-seq data. The authors performed RNA-seq analyses with female gametocytes collected by cell sorting using a female-specific fluorescent marker and compared the female transcriptome of *P. yoelii* 17XNL (*Py*WT) and *pyap2-fg2*-null parasites. The analysis detected significant upregulation of 1141 genes in *pyap2-fg2*-null parasites, more than half of which were specifically or preferentially expressed in males. Accordingly, the authors concluded that *Py*AP2-FG2 globally represses male genes to safeguard the female transcriptome. This statement differed from our conclusion that *Pb*AP2-FG2 and AP2R-2 repress various genes, which include some early gametocyte and female genes. The second is the conclusions derived from their ChIP-seq analysis of *Py*AP2-FG2. For example, the binding motif of *Py*AP2-FG2 predicted in their study was considerably different from that of *Pb*AP2-FG2 identified in this study; the binding motif of *Py*AP2-FG2 was predicted to be TRTRTGCA. As *P. berghei* and *P. yoelii* are phylogenetically very close, such discrepancies in the roles of orthologous genes seems implausible. Therefore, we reassessed the ChIP-seq and RNA-seq data deposited from the *Py*AP2-FG2 study to clarify the inconsistency between the two studies.

First, we analyzed the ChIP-seq data for *Py*AP2-FG2. We mapped their sequence data onto the *P. yoelii* reference genome (downloaded from PlasmoDB) using bowtie2 and removed reads aligned onto more than two sites of the genome, as in our ChIP-seq analysis. Then, we called peaks using macs2 with the criteria used by Li *et al*. (fold enrichment > 2.0 and *p*-value < 1.0 × 10^-5^) and obtained 1309 peaks that were common in the duplicates (S6A and S6B Table). Within 100 bp from these peak summits, we found enrichment of TRTRTGCA with a *p*-value of 7.0 × 10^-52^ by Fisher’s exact test (Fig. 8A). On the other hand, the major motif identified in our study, RGAGAR, was also enriched with a *p*-value of 2.2 × 10^-52^, comparable to that of the TRTRTGCA motif (Fig 8A). In addition, the variant motifs 1 and 2 were also enriched with *p*-values of 4.0 × 10^-48^ and 9.4 × 10^-26^, respectively. We further performed IDR1D analysis and found that only 122 peaks had an IDR < 0.01, which implies the low reproducibility of the ChIP-seq data (Fig 8B and 8C, S6A and S6B Table). Within these highly reliable peaks (IDR < 0.01), the RGAGAR motif was much more enriched (*p*-value = 2.1 × 10^-12^) than the TRTRTGCA motif (*p*-value = 1.1 × 10^-4^, Fig. 8A). Therefore, in contrast to the major motif, the TRTRTGCA motif appeared to have been mainly derived from unreliable peaks (IDR ≥ 0.01).

**Fig 8.**
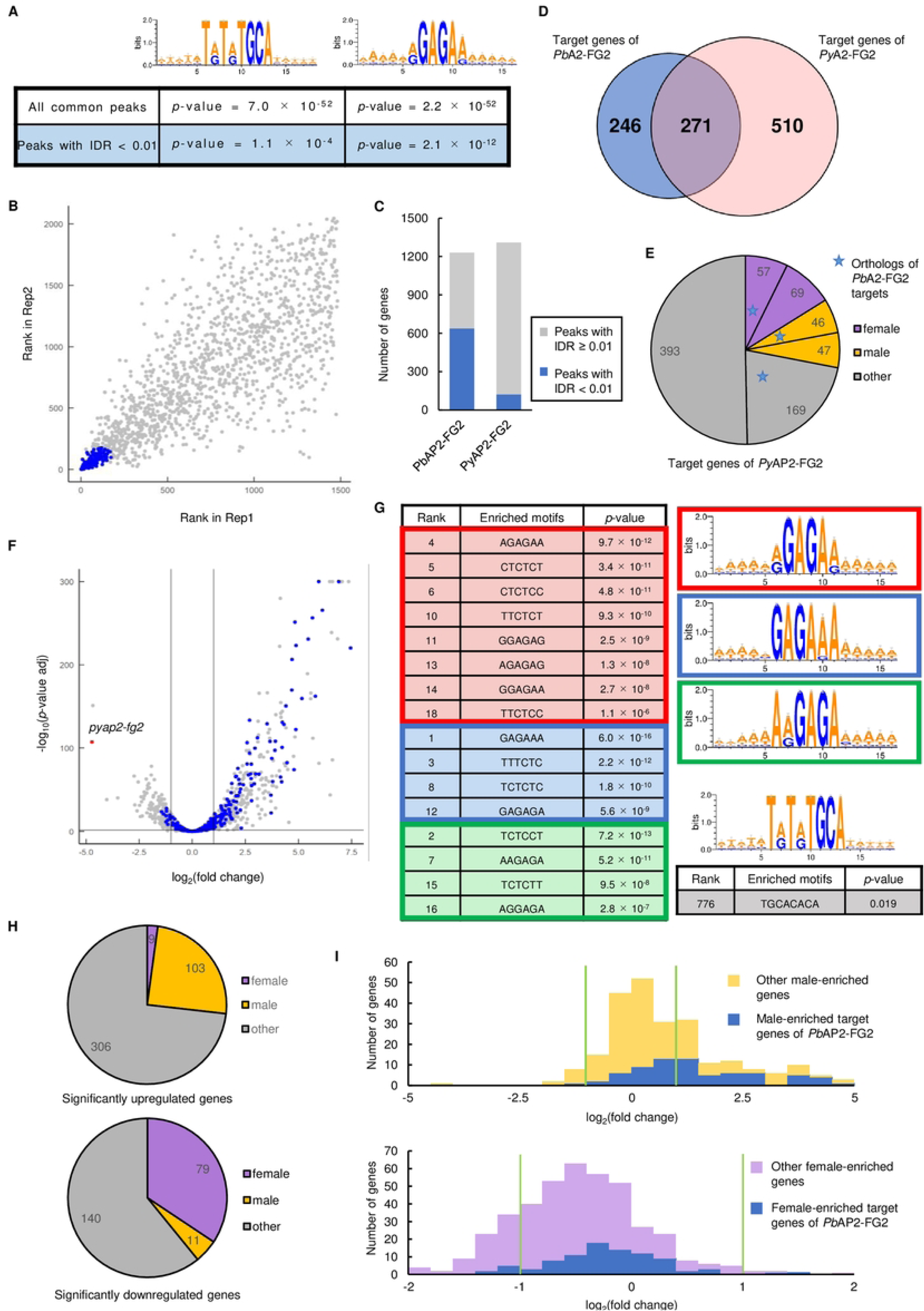
Reassessment of the ChIP-seq and RNA-seq data for the study of *Py*AP2-FG2. (A) Enrichment of TRTRTGCA and RGAGAR motif in the peak region identified by ChIP-seq analysis of *Py*AP2-FG2. The logos were depicted by WebLogo 3. (B) IDR1D analysis between the ChIP-seq experiment 1 and 2 of *Py*AP2-FG2. Peaks with IDR < 0.01 are indicated as blue dots. (C) Ratio of peaks with IDR < 0.01 in all peaks identified in ChIP-seq of *Pb*AP2-FG2 and *Py*AP2-FG2. (D) A Venn diagram showing number of genes common in the targets of *Pb*AP2-FG2 and *Py*AP2-FG2. (E) Classification of the target genes of *Py*AP2-FG2 into sexual stage-enriched gene sets. The number of target genes common for *Py*AP2-FG2 and *Pb*AP2-FG2 are indicated with a blue star for each set. (F) A volcano plot showing DEGs in *pyap2-fg2*-null parasite compared to *Py*WT. Blue dots represent orthologs of the target genes of *Pb*AP2-FG2, and a red dot indicates *pyap2-fg2*. A horizontal line indicates a *p*-value of 0.05, and two vertical lines indicate log_2_(Fold Change) of -1 and 1. (G) Six-bp DNA motifs enriched within the upstream region (300 to 1200 bp from ATG) of genes upregulated in *pyap2-fg2*-null parasite. (H) Classification of significantly upregulated and downregulated genes into sexual stage-enriched gene sets (top and bottom, respectively). (I) Histograms showing distribution of log_2_(Fold Change) values in *pyap2-fg2*-null parasite for female and male-enriched genes (top and bottom, respectively). Green lines indicate log_2_(Fold Change) of -1 and 1.

We further predicted the target genes of *Py*AP2-FG2 from the 1309 peaks common in duplicates. The analysis revealed that 781 target genes, of which 271 orthologous genes in *P. berghei*, were also identified as the target genes of *Pb*AP2-FG2 (Fig 8D and S6C Table). Although there was a significant overlap between the two target sets (*p*-value = 2.4 × 10^-85^ by Fisher’s exact test), approximately 70% of the *Py*AP2-FG2 targets were not included in the *Pb*AP2-FG2 targets. Given the low reproducibility of the ChIP-seq data, we supposed that some targets might have been falsely detected. We further assessed the sex-specific expression of the target genes of *Py*AP2-FG2 and found that they contained only 94 male-enriched genes. This result contradicts the conclusion that *Py*AP2-FG2 globally represses male genes (Fig 8E). Moreover, the targets of *Py*AP2-FG2 also contained 128 female-enriched genes, 58 orthologous genes of which were included in the targets of *Pb*AP2-FG2, implying that, similar to *Pb*AP2-FG2, *Py*AP2-FG2 also plays a role in repressing a substantial number of female genes.

Next, we assessed the RNA-seq data of *the Py*WT and *pyap2-fg2*-null parasites. In their RNA-seq analysis, no threshold of fragments per kilobase of transcript per million mapped reads (FPKM) was set to exclude genes with low expression levels. Such an analytical process may detect DEGs derived from artificial variances and yield unreliable DEG lists. In an analysis performed by Li *et al*., more than one-third of the upregulated genes (445/1,141 genes) had FPKM < 10 for *pyap2-fg2*-null parasites. Although these genes with low scores could demonstrate high-fold enrichment, the actual upregulation was low and could be false positives; thus, it is not appropriate to conclude the function of *Py*AP2-FG2 from such analyses. To obtain more robust results, we analyzed the RNA-seq data according to the analytical process performed in this study, setting a minimum FPKM threshold of < 10 (S7A Table). This analysis identified 418 significantly upregulated and 230 significantly downregulated genes (Fig 8F, S7B and S7C Table). In the upregulated genes, the target genes of *Py*AP2-FG2, which we obtained above, were enriched with a *p*-value of 7.7 × 10^-9^ by Fisher’s exact test. On the other hand, the orthologous genes of *Pb*AP2-FG2 targets were more enriched (*p*-value = 3.4 × 10^-17^) (Fig 8F), again suggesting that the target list of *Py*AP2-FG2 may contain several pseudo-targets. We next investigated whether the binding motifs of AP2-FG2 identified in this study and that by Li *et al*. were enriched in the upstream of these upregulated genes. Consistent with our results, the major motif and two variant motifs were highly enriched in the upstream region compared to that of the other genes (Fig 8G). In fact, all 6-bp motifs that belonged to the major motif, variant motif 1 or 2 were detected in the 20 most enriched motifs, suggesting that these motifs functioned as a *cis*-acting repressive element in *P. yoelii* as well. However, when enrichment of any 8-bp motif was searched in the same region, the TRTRTGCA motif was not found to be significantly enriched; that is, the most enriched motif that corresponds to TRTRTGCA was detected as the 776th enriched motif with a *p*-value of 0.019 (Fig 8G), indicating that this motif is not related to the upregulation of genes detected in *pyap2-fg2*-null parasites.

According to our analysis, majority of male-enriched genes were not significantly upregulated in *pyap2-fg2*-null parasites; the upregulated genes only contained 103 male-enriched genes (Fig 8H and 8I). Moreover, nearly half of the upregulated male-enriched genes were not a target gene of *Pb*AP2-FG2 or *Py*AP2-FG2. Therefore, based on our analysis of their RNA-seq data, it seemed not appropriate to conclude that *Py*AP2-FG2 globally represses male genes. For female-enriched genes, only nine genes were significantly upregulated, and most genes tended to be downregulated, as discussed by Li *et al*. (Fig 8H and 8I). The downregulation of female genes was considered to be caused by impairment of the female transcriptome upon disruption of *pyap2-fg2*. Therefore, we hypothesized that such an effect caused by the disruption of *pyap2-fg2* might have masked the upregulation of female-enriched target genes in the RNA-seq analysis. In fact, despite the overall downregulation of female-enriched genes, most of the female-enriched target genes were not downregulated; there was a significant difference in the distribution of log_2_(fold change) between the female-enriched target genes and the other female-enriched genes (*p*-value = 1.7 × 10^-4^ by two-tailed Student’s t-test) (Fig 8I). These results suggested that consistent with the results for *Pb*AP2-FG2, *Py*AP2-FG2 also targets female genes and represses their expression. Collectively, our analysis revealed that *Pb*AP2-FG2 and *Py*AP2-FG2 both repress not only male genes but also a wide-variety of genes to support female differentiation.

## Discussion

This study highlights how *Pb*AP2-FG2 and AP2R-2 function cooperatively as a transcriptional repressor complex during female development. Their target genes contained variable genes regarding functional annotation and expression patterns, which indicated that this repressor complex might play distinct roles for each group of the target genes during female development. As suggested by the comparison between the target genes of *Pb*AP2-FG2 and AP2-G, one of the roles of the repressor complex could be repression of the early gametocyte genes to promote female differentiation. During early gametocyte development, the female transcriptional activator AP2-FG begins to be expressed as AP2-G expression decreases [11, 14]. Thus, we considered that the repression of AP2-G targets was vital for completing this switch of major transcriptional activators. On the other hand, the target genes of *Pb*AP2-FG2 also included a significant number of female-enriched genes. This result appeared unreasonable because it would mean that such genes are activated and repressed during the same period. Nevertheless, the differential expression analysis suggested that these female-enriched target genes were indeed repressed by *ap2-fg2*; for both *P. berghei* and *P. yoelii*, expression of the female-enriched target genes was not downregulated in *ap2-fg2*-knockout parasites despite that the other female-enriched genes were predominantly downregulated. This observation implied that transcriptional activators alone could not precisely control gene expression for female gametocyte development, thereby requiring repressors for its modulation. Another possible role of the *Pb*AP2-FG2-AP2R-2 repressor complex was the repression of male genes, as suggested by Li et al. [18]. However, the results of this study did not corroborate their conclusion that the global repression of male genes by *Py*AP2-FG2 was required for balancing the female-specific transcriptome. The ChIP-seq analyses of AP2-FG2 in *P. berghei* and *P. yoelii* both showed that the target genes of AP2-FG2 contained only a subset of male-enriched genes. These included only a limited number of genes related to the major characteristic features of male gametocytes/microgametes, such as flagella formation and DNA replication. Moreover, although a significant number of male-enriched genes were upregulated in *pyap2-fg2*-null parasites, nearly half of them were not a target gene of AP2-FG2. From this result, we speculated that the target genes of AP2-FG2 could contain male-specific transcriptional regulator genes, especially activators; the non-target genes were indirectly upregulated in *pyap2-fg2*-null parasites, as the expression of male transcriptional activators was released from the repression by *Py*AP2-FG2. Therefore, we proposed that the repressor complex repressed a subset of male genes that include those essential for regulating male differentiation. Collectively, our results helped us conclude that the AP2-FG2-AP2R-2 complex could play three roles: promoting female differentiation by repressing early gametocyte genes, modulating the expression level of female genes, and suppressing male differentiation.

It has been reported that *Plasmodium* parasites possess another transcriptional repressor, AP2-G2 [12, 13]. *ap2-g2* is activated by AP2-G [11], a transcriptional activator that triggers sexual differentiation, similar to *pbap2-fg2*, which is activated by the female transcriptional activator AP2-FG. AP2-G2 induces global gene repression, and disruption of this gene results in the aberrant development of gametocytes. The target genes of AP2-G2 include various genes, like those of *Pb*AP2-FG2 do, suggesting that AP2-G2 also plays multiple roles, possibly repressing trophozoite genes, modulating the expression of early gametocyte genes, and suppressing asexual fate. Studies of *Plasmodium* AP2-family proteins have shown that transcriptional regulation by AP2 transcription factors is remarkably simple; one transcription factor comprehensively activates certain stage-specific genes [11,14,15,36,37]. On the other hand, recent studies of repressors have indicated that to regulate stage conversion, such as sexual differentiation and sex determination, transcriptional repressors also play an essential role [12,13,18,38]. In addition, studies in another Apicomplexan parasite, *Toxoplasma*, have reported an essential role for transcriptional repressors in conversion from tachyzoite to bradyzoite [39, 40]. Therefore, it was suggested that optimal transcriptional regulation could not to be controlled by transcriptional activators alone during stage conversion in Apicomplexa. This observation suggested the possibility of finding transcriptional repressors during the other stage conversion processes as well, and investigating these factors would help us understand the mechanisms promoting the life cycle of *Plasmodium*.

## Materials and Methods

### Ethical statement

All experiments in this study were performed following the recommendations in the Guide for the Care and Use of Laboratory Animals of the National Institutes of Health to minimize animal suffering and were approved by the Animal Research Ethics Committee of Mie University, Mie, Japan (permit number 23–29).

### Parasite preparation

*pbap2-fg2*(-), *Pb*AP2-FG2::GFP, and *ap2r-2*(-) were derived from the *P. berghei* ANKA strain. The other transgenic parasites were derived from Pbcas9 [28]. Parasites were inoculated in Balb/c or ddY mice. Ookinete cultures were performed at 20 °C using RPMI1640 medium, pH 8.0, supplemented with fetal calf serum and penicillin/streptomycin at final concentrations of 20% and 100 U/mL, respectively. Cross-fertilization assays were conducted as previously described [41].

### Generation of mutant parasites

The DNA constructs for tagging *Pb*AP2-FG2 with GFP and knocking out *pbap2-fg2* were prepared as previously reported [41, 42]. Briefly, for *gfp*-tagging, two homologous regions were cloned into the *gfp*-fusion vector to fuse *pbap2-fg2* in-frame with *gfp*. This vector possesses a *hdhfr* expression cassette next to the *gfp* so that mutants can be selected by treatment with pyrimethamine. The plasmid was linearized by *Xho*I and *Not*I digestion before use in transfection experiments. To knock out *pbap2-fg2*, the targeting construct was prepared using overlap PCR. The construct had two homologous regions around the *pbap2-fg2* locus flanking a *hdhfr* expression cassette. *ap2r-2*(-) was generated in a previous study [11].

The other transgenic parasites were generated by the CRISPR/Cas9 system using the parasites expressing Cas9 [28]. The Cas9-expressing parasite Pbcas9 has a *cas9* cassette at the *p230p* locus. The *hsp70* promoter controls the expression of Cas9, and Pbcas9 constitutively expresses Cas9 throughout the asexual blood cycle. Donor DNA for transfection was constructed by overlap PCR, cloned into pBluescript KS (+) using the *Xho*I and *Bam*HI sites by In-Fusion cloning, and then amplified by PCR from the constructed plasmid. sgRNA vectors were constructed as previously described [28]. Target sites of sgRNA were designed using the online tool CHOPCHOP (https://chopchop.cbu.uib.no/).

Transfection was performed using the Amaxa Basic Parasite Nucleofector Kit 2 (LONZA). All transfectants were selected by treating mice with 70 μg/mL pyrimethamine in their drinking water. Recombination was confirmed by PCR and/or Sanger sequencing, and clonal parasites were obtained by limiting dilution. All primers used in this study are listed in S8 Table (No. 1–48).

### RNA-seq and sequence data analysis

The total RNA was extracted from parasites enriched with gametocytes by treating infected mice with 10 mg/L sulfadiazine in their drinking water for two days, using the Isogen II reagent (Nippon gene). Briefly, whole blood was withdrawn from infected mice and passed through the Plasmodipur filter to remove white blood cells, and the red blood cells (RBCs) were lysed in an ice-cold 1.5 M NH_4_Cl solution. After the lysis, the cells were subjected to Isogen II (NIPPON GENE), and the total RNA was extracted according to the manufacturer’s instructions. RNA-seq libraries were prepared from the total RNA using the KAPA mRNA HyperPrep Kit (Kapa Biosystems) and sequenced using Illumina NextSeq. Three biologically independent experiments were conducted for each parasite line. The obtained sequence data were mapped onto the reference genome sequence of *P. berghei*, downloaded from PlasmoDB 46, using HISAT2, setting the parameter for maximum intron length to 1000. The mapping data for each sample were analyzed using featureCounts and compared using DESeq2. Genes in the subtelomeric regions were removed from the differential expression analysis. The parameters for all programs were set as the default unless otherwise specified.

### ChIP-seq and sequencing data analysis

Whole blood was withdrawn from the infected mice treated with sulfadiazine and passed through the Plasmodipur filter to remove white blood cells. The blood was diluted in a complete medium (RPMI1640 supplemented with 20% fetal calf serum) and immediately fixed with 1% formalin at 30 °C. After fixing, RBCs were lysed in ice-cold 1.5 M NH4Cl solution. This step was performed several times until the supernatant became clear, and the cells were lysed in SDS lysis buffer. The samples were sonicated using Bioruptor (Cosmo Bio) for 20 cycles of 30 sec on/30 sec off to shear the chromatin. Chromatins were immunoprecipitated with anti-GFP polyclonal antibodies (Abcam), which were bound to Protein A Magnetic Beads (Invitrogen) before the ChIP step. DNA fragments were purified from the immunoprecipitated chromatin and used for library construction. Libraries for NGS were prepared using the KAPA HyperPrep Kit (Kapa Biosystems) according to the manufacturer’s instructions and sequenced using Illumina NextSeq. Two biologically independent experiments were performed for each sample and used for the following analysis.

The obtained sequence data were mapped onto the reference genome sequence of *P. berghei* using Bowtie 2. Reads aligned onto more than two sites were removed from the mapping data. Using the trimmed mapping data, peaks were called with macs2 callpeak with fold enrichment > 3.0 and *q*-value < 0.01. To identify reliable peaks, the data obtained from two biologically independent experiments were compared using IDR1D analysis (https://idr2d.mit.edu/) setting max gap to 100. Briefly, the peaks were ranked for each replicate according to their *p*-value. The peaks from each replicate were then compared and scored based on their respective ranks. Highly reproducible peaks were defined as those with an IDR score < 0.01. Binding motifs were predicted by analyzing the enrichment of motifs within 50 bp of peak summits using Fisher’s exact test (the method was previously described in detail [36]). Genes with peaks within upstream of 1200 bp from ATG were identified as target genes. The parameters for all programs were set as the default unless otherwise specified.

### ChIP-qPCR and RT-qPCR for reporter experiments

ChIP experiments were performed as described for ChIP-seq analysis. Quantification of DNA fragments of interest was performed by real-time qPCR using TB Green Fast qPCR Mix (Takara) and Thermal Cycler Dice Real Time System II (Takara). Three biologically independent experiments were performed and used for the analysis.

For RT-qPCR analyses, cDNA was synthesized from total RNA, extracted as described for RNA-seq analysis, using the PrimeScript RT reagent Kit with gDNA Eraser (Takara). Real-time qPCR experiments were performed as described above. All primers used are listed in Table S8 (No. 49–62).

## Figure captions

### Supporting information

**S1 Fig. Genotyping of transgenic parasites developed in this study.** (A) *Pb*AP2-FG2::GFP. (B) *pbap2-fg2*(-). (C) *Pb*AP2-FG2::GFP^C^. (D) AP2R-2::GFP^C^.

**S1 Table. List of differentially expressed genes in *pbap2-fg2*(-).** (A) RPKM values in each data. (B) Significantly downregulated genes. (C) Significantly upregulated genes.

**S2 Table. List of sex-enriched genes.** (A) Female-enriched genes. (B) Male-enriched genes.

**S3 Table. List of peaks and target genes identified in the ChIP-seq experiments of *Pb*AP2-FG2.** (A) Peaks in Experiment 1. (B) Peaks in Experiment 2. (C) Target genes.

**S4 Table. List of peaks identified in the ChIP-seq experiments of AP2R-2.** (A) Peaks in Experiment 1. (B) Peaks in Experiment 2.

**S5 Table. List of differentially expressed genes in *ap2r-2*(-).** (A) RPKM values in each data. (B) Significantly downregulated genes. (C) Significantly upregulated genes.

**S6 Table. List of peaks and target genes identified in the ChIP-seq experiments of *Py*AP2-FG2.** (A) Peaks in Experiment 1. (B) Peaks in Experiment 2. (C) Target genes.

**S7 Table. List of differentially expressed genes in *pyap2-fg2*-null parasite.** (A) FPKM values in each data. (B) Significantly downregulated genes. (C) Significantly upregulated genes.

**S8 Table. List of primers used in this study.**

## Notes

### Competing Interest Statement

The authors have declared no competing interest.

